# Single Cell Atlas of Human Putamen Reveals Disease Specific Changes in Synucleinopathies: Parkinson’s Disease and Multiple System Atrophy

**DOI:** 10.1101/2021.05.06.442950

**Authors:** Rahul Pande, Yinyin Huang, Erin Teeple, Pooja Joshi, Amilcar Flores-Morales, Martine Latta-Mahieu, S. Pablo Sardi, Angel Cedazo-Minguez, Katherine W. Klinger, Stephen L. Madden, Deepak Rajpal, Dinesh Kumar

## Abstract

Understanding disease biology at a cellular level from disease specific tissues is imperative for effective drug development for complex neurodegenerative diseases. We profiled 87,086 nuclei from putamen tissue of healthy controls, Parkinson’s Disease (PD), and Multiple System Atrophy (MSA) subjects to construct a comprehensive single cell atlas. Although both PD and MSA are manifestations of α-synuclein protein aggregation, we observed that both the diseases have distinct cell-type specific changes. We see a possible expansion and activation of microglia and astrocytes in PD compared to MSA and controls. Contrary to PD microglia, we found absence of upregulated unfolded protein response in MSA microglia compared to controls. Differentially expressed genes in major cell types are enriched for genes associated with PD-GWAS loci. We found altered expression of major neurodegeneration associated genes — SNCA, MAPT, LRRK2, and APP — at cell-type resolution. We also identified disease associated gene modules using a network biology approach. Overall, this study creates an interactive atlas from synucleinopathies and provides major cell-type specific disease insights.

Link to interactive atlas will be made available at the time of publication.

## Introduction

Synucleinopathies consist of a group of neurodegenerative disorders characterized by the presence of α-synuclein protein aggregates in the brain. These disorders include Parkinson’s disease (PD), Alzheimer’s Disease (AD), dementia with Lewy bodies (DLB), pure autonomic failure (PAF) and Multiple System Atrophy (MSA) [1]. Parkinson’s disease is the second most frequent neurodegenerative disorder, next to Alzheimer’s. However, MSA is considered a rare disease with an estimated incidence of 0.6-3 per 100,000 people [2]. Although α-synuclein aggregates are a hallmark of the Synucleinopathies, recent studies have shown the presence of Lewy-related pathology (LRP), mostly comprised of α-synuclein in more than 50% of autopsy-confirmed AD brains [3, 4], implicating α-synuclein accumulation in the pathophysiology of Alzheimer’s [5, 6]. These neurodegenerative diseases often exhibit overlapping clinical symptoms which makes differential early diagnosis challenging. Clinically, Synucleinopathies exhibit chronic progressive decline in motor, cognitive, behavioral, and autonomic function [1].

Despite significant advances in our understanding of Synucleinopathies, there remains a lot to be understood about the cellular and molecular underpinnings of these diseases, including the α-synuclein cellular biology. Recent developments in single cell techniques have allowed comprehensive molecular profiling of human brain tissue at single cell resolution. To date, we are aware of a lone single-cell study from human PD midbrain [7] and none from MSA. We believe that this is the first comprehensive study to the best of our knowledge investigating two different synucleinopathies at the single cell level. We present a single-nucleus atlas of the representative synucleinopathies, PD and MSA by single cell RNA sequencing of a total of 87,086 nuclei from healthy controls, PD samples, and MSA samples (Control – 22,297; MSA – 32,488; PD – 32,301). The data will be available in the form of a web portal with interactive access :

(link will be made available at the time of publication)

While PD and MSA are both neurodegenerative diseases with several shared manifestations, we suggest, based on this study data that there are significant differences at the molecular level which could potentially influence selective therapeutic development efforts for PD and MSA. We observed trends of altered proportions of OPC, astrocytes and microglia in disease condition as compared to controls. To investigate the disease specific states in different cell types, we performed trajectory reconstruction using Monocle2 [8] and observed disease specific branching path in microglia. We identified differentially expressed genes between disease and controls within the major brain tissue cell subtypes and the possible biological pathways affected by these changes. These findings point to distinct molecular signatures at the subcellular level, that accompany pathological diagnoses of these diseases and can help in improving diagnostic methods. Differentially expressed genes for PD cell types had statistically significant overlap with gene loci nearest to PD GWAS variants. The expression of genes associated with PD, SNCA and LRRK2, were found to be overexpressed in PD microglia. Moreover, the expression of MAPT, associated with several neurodegenerative diseases, was upregulated in PD oligodendrocytes and OPCs. By performing WGCNA, we identified a common gene module implicated in both PD and MSA which could have potential applications in drug discovery. This atlas of Synucleinopathies of human putamen will accelerate the elucidation of α-synuclein biology in a disease specific manner at the cellular level. We believe that these data will serve as a valued resource in uncovering novel biology in PD and MSA pathogenesis.

## Results

### Alteration of Cell-Type Heterogeneity in PD and MSA

The human brain is heterogenous in terms of cell types and region-specific differences are not known for many of the brain regions including putamen in the midbrain [9]. Therefore, as a first step we wanted to establish the baseline proportion of the major cell types and then investigate how their relative proportions change within PD and MSA. We performed single nuclei sequencing of postmortem human putamen tissue samples from nine subjects (3 Control, 3 clinically diagnosed with PD, 3 clinically diagnosed with MSA) as illustrated in Figure 1A. After quality control filtering, as described in the Methods section, we retained 22,297 Control nuclei, 32,301 PD nuclei and 32,488 MSA nuclei. We integrated data from all samples to align cell types across Control, PD and MSA, and subsequently performed nuclei clustering. The derived cell clusters were then classified, annotated for major cell types using canonical markers, (Supplementary Table 1) and quantified. We found the presence of all major brain cell-types in the putamen, ordered in decreasing proportion: Oligodendrocytes (62.88%), Neurons (17.42%), Astrocytes (9.98%), Microglia (5.21%), OPCs (Oligodendrocyte Precursor Cells) (3.76%), and VLMC (Vascular and Leptomeningeal Cells) (0.72%). Figure 1B shows the t-SNE plot of nuclei and expression of marker genes across different cell types. No statistically significant difference in the proportion of the different cell types was established between the three groups, although we observed an increasing trend in astrocytes and microglia proportion, and decreased proportion of OPC in PD as compared to Control tissues (Figure 1C). Similarly, we also observed a reduced astrocyte proportion in MSA as compared to Control tissues (Figure 1C). The VLMC make up < 1% of total nuclei, so we do not speculate on the change in VLMC proportion.

**Figure 1.**
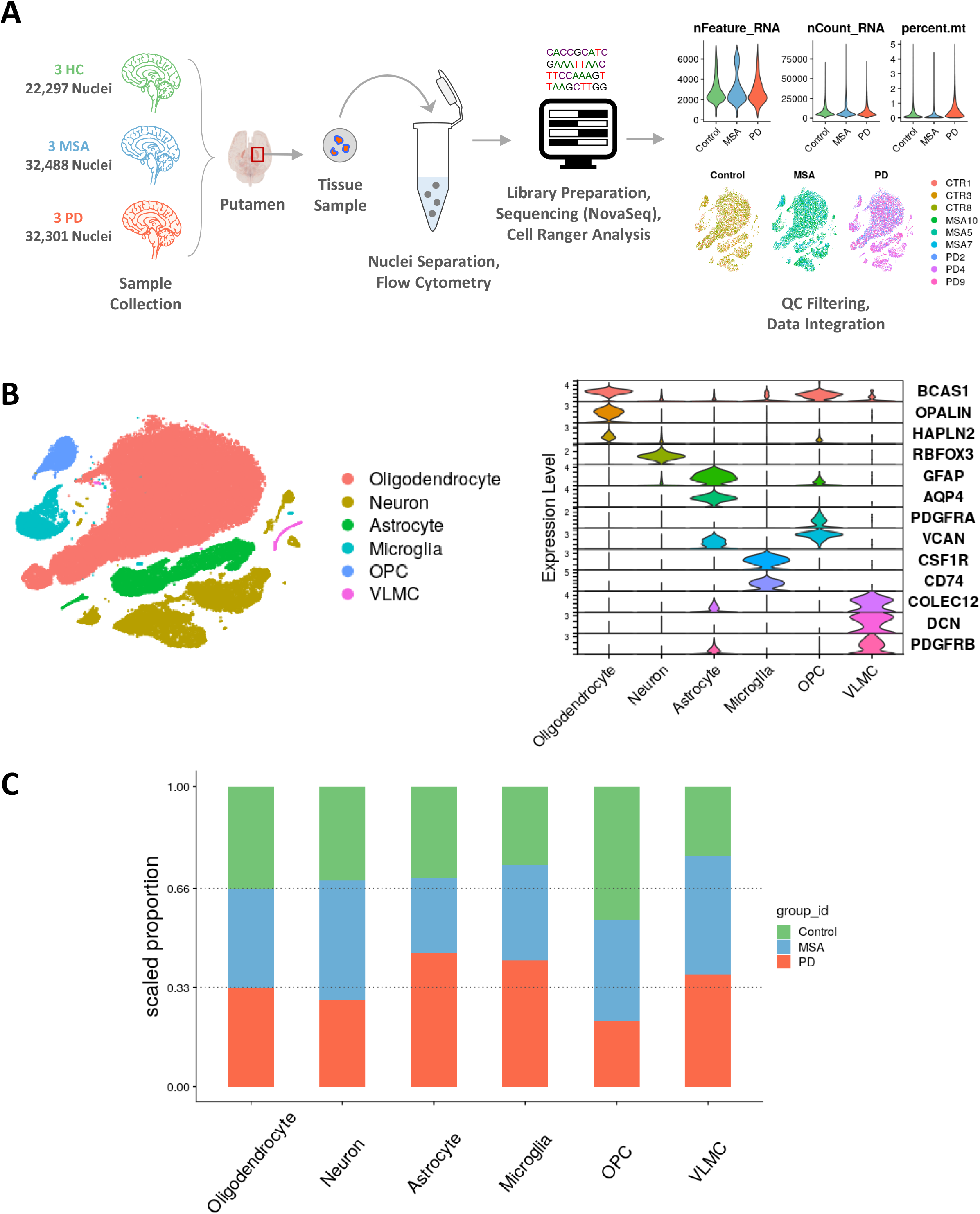
Broad Cell Type Identification. **(A)** Schematic workflow outlining samples processing, library preparation, quality control and subsequent sample integration **(B)** Gene expression profile of nuclei clusters established presence of major brain cell types **(C)**Trends of altering OPC, astrocytic and microglial proportion in PD and MSA compared to Control

### Disease Specific Changes in PD and MSA Neurons

In order to study the neuronal population in detail, we re-clustered the neuron subset, as described in Methods section. The resulting clusters were annotated based on differentially expressed genes across clusters in the Control group and by previously identified marker genes. Expression of PPP1R1B (also known as DARPP-32) is characteristic of striatal medium spiny projection neurons (SPNs) [10], which was observed as the most abundant neuronal cell type in putamen (93.07%). We identified five types of neurons; PPP1R1B^+^/DRD1^+^ medium spiny projection neurons (D1MSN); PPP1R1B^+^/DRD2^+^ medium spiny projection neurons (D2MSN); a distinct spiny neuronal population (MSN) characterized by expression of PPP1R1B^+^/ADARB2^+^; and two GAD1^high^/GAD2^high^ GABAergic inhibitory neuronal clusters (GABA1 and GABA2), as shown in Figure 2A. We did not find any SLC17A7^+^ glutamatergic/excitatory neurons, which are abundant in cortical areas [11].

**Figure 2.**
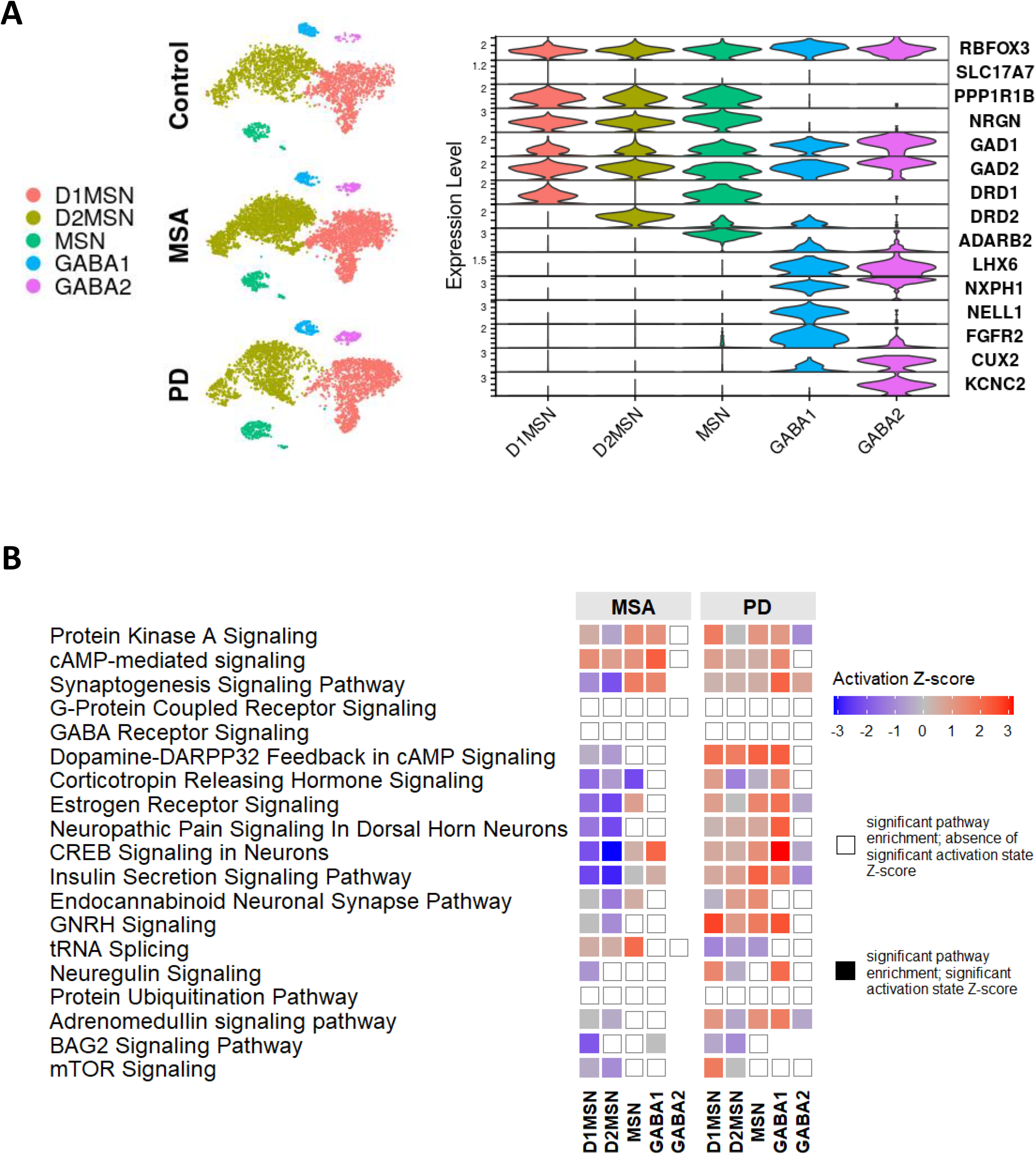
Neuronal Heterogeneity. **(A)** Molecular profiling reveals major neuronal phenotypes -D1 receptor spiny projection neurons (D1MSN); D2 receptor spiny projection neurons (D2MSN); GABAergic neurons (GABA1 and GABA2) and an unresolved ADARB2+ /PPP1R1B+ (MSN) cluster **(B)** Shows contrasting regulation of enriched biological pathways in MSA and PD neuronal subtypes compared to Control group; Color signifies pathway activation zscore, assigned by Ingenuity Pathway Analysis (IPA) software, whereas unfilled marks represent enriched pathways with no available zscore from IPA

For each of the disease state neuronal sub-types, the differentially expressed genes, compared to Control, were analyzed for biological pathway enrichment using IPA. Figure 2B shows differential regulation of pathways within neuronal sub-types for PD versus Control and MSA versus Control, where color scale represents the relative activation z score of the enriched pathway. We found contrasting changes in the pathways that are seen to be modulated particularly for D1MSN and D2MSN neurons in PD and MSA. The transcription pattern of genes related to CREB signaling, Ca+ signaling, synaptic transmission and dopamine feedback was downregulated in MSA neuronal subtypes, whereas in contrast were relatively upregulated in the case of PD (Figure 2B). Several genes involved in calcium transport (CACNA1I, CACNG2, CACNA2D3), cytoplasmic phospholipase gene - PLCG2 and G-Protein coupled receptor genes GPR156, GPR137C were downregulated in MSA D1/D2MSN neurons, whereas, in the case of PD D1/D2MSN neurons, these genes were upregulated. Expression of the anti-apoptotic gene BCL2 was downregulated in MSA D1/D2MSN neurons. Growth factor genes GHR, TGFB2, TGFBR3 were downregulated in both PD and MSA spiny neurons compared to Control. A comprehensive list of all differentially expressed genes for all neuronal sub types is provided in Supplementary Table 2.

### Gene Co-Expression Network Analysis Reveals Disease Correlated Gene Set in Neurons

To gain more insight into the disease biology and to identify disease associated transcriptional changes, we performed Weighted Gene Co-expression Network Analysis (WGCNA) for neurons and obtained 12 gene co-expression modules as shown in Figure 3A. Pearson correlations of module eigen genes with one-hot encoded disease condition is shown in Figure 3B. Interestingly, we noticed that the Brown module had the strongest association with PD as well as MSA (p values < 1e-323). The corresponding Brown module eigen gene had strong positive correlation (0.72) with MSA and a strong negative correlation (−0.7) with PD (p value < 1e-323 for both associations). The Brown module contains 417 genes including genes DDX5, SPRED1, ATR involved in DNA damage response. It also consists of mRNA processing/splicing related genes such as RBPMS, RNMT, RNPC3, PDCD7, PRPF4B, ELAVL1, PNN, GCFC2, SON, SRSF 7/10/11 and SF1. To aid in the biological interpretation of this module, we built a Protein-Protein functional Interaction (PPI) network using Brown module genes as seeds (Figure 3C). Analysis of this network revealed ELAVL1, FN1, DDX5, TUBA1A, CFTR, APP as the top hub genes (Figure 3C). Amyloid Beta Precursor Protein (APP) has been previously linked with neurodegenerative diseases [12]. KEGG Pathway enrichment of the network proteins (Figure 3D) implicates PD and MSA associated pathways linked to apoptosis, autophagy, dopamine feedback and proteolysis, among others. The PPI network had 14 genes in common with genes within 100kb up and 100kb down promoter regions of PD GWAS variants (association p value < 1e-5) -UBE3A, UBQLN4, FUS, SHC1, BAG3, MKRN3, CCT3, TTC19, CLK2, KCNH2, CDK5, DVL3, LRRK2, and SPDYA (Figure 3E).

**Figure 3.**
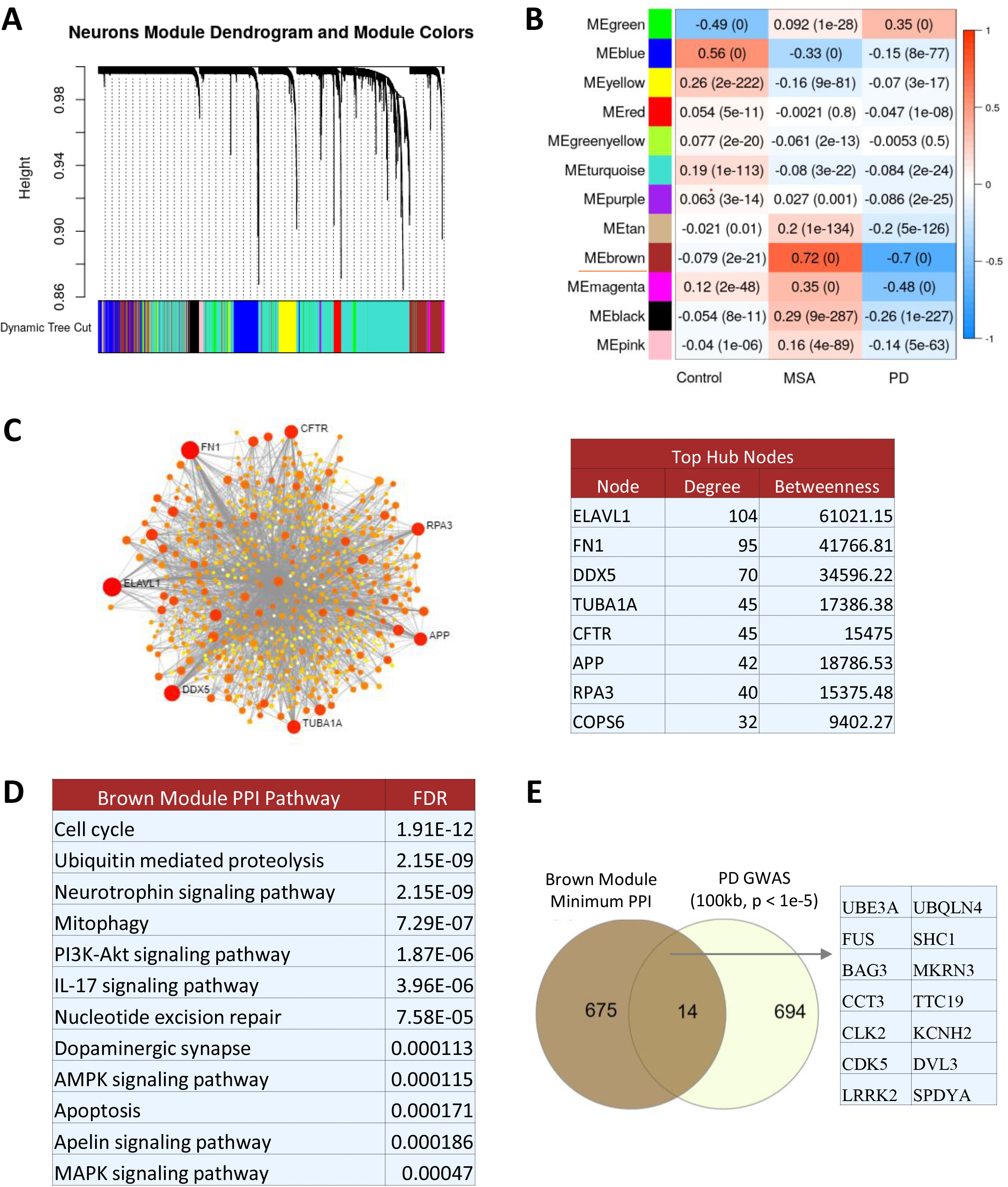
Gene Co-Expression Analysis. **(A)**. Gene co-expression clustering dendrogram based on topological overlap dissimilarity **(B)** Brown module is significantly associated with both PD and MSA **(C)** Functional Protein-Protein Interaction (PPI) network of Brown module genes reveals APP as one of the hub nodes **(D)** Brown PI network genes are enriched for dopaminergic synapse, mitophagy and apoptosis pathways **(E)** Brown PI network genes have 14 overlapping genes with PD GWAS (100kb region of SNPs with association p<1e-5)

### PD and MSA Microglia Have Distinct Changes in the Disease Related Pathways

Microglia are the major immune cells in the brain and have been shown to play a critical role in neurodegenerative biology [13]. To investigate microglial population in detail, we partitioned and re-clustered microglia for further analysis, considering the first 10 principal components and a 0.1 resolution parameter (Figure 4A). Two clusters (MG1 and MG2) expressed canonical microglial markers TYROBP, P2RY12, AND CSF1R and the other two clusters were identified as peripheral cell types, macrophages, and T cells. The macrophage cluster was distinguished by expression of MRC1 and TLR4 and T-cell cluster with IL7R and CCL5 expression. IPA pathway enrichment from results of differential gene expression of PD versus Control and MSA versus Control clusters, is shown in Figure 4B. The Unfolded Protein Response (UPR) pathway was upregulated in both PD-MG1 and PD-MG2 nuclei compared to Control. We also observed over expression of several heat shock proteins HSPH1, HSP90AA1, HSP90AB1, HSPA1A/B, HSP90B, HSPA8/6/4 (members of UPR pathway) in PD-MG1 and PD-MG2 clusters compared to Controls. MSA microglia were enriched for antigen presentation and neuroinflammation pathway, with upregulation of chemokine ligand CXCL2 and Major Histocompatibility Complexes HLA-DRB1, HLA-DPA1 and HLA-C genes. Please refer to Supplementary Table 3 for extensive list of differentially expressed genes for microglia clusters.

**Figure 4.**
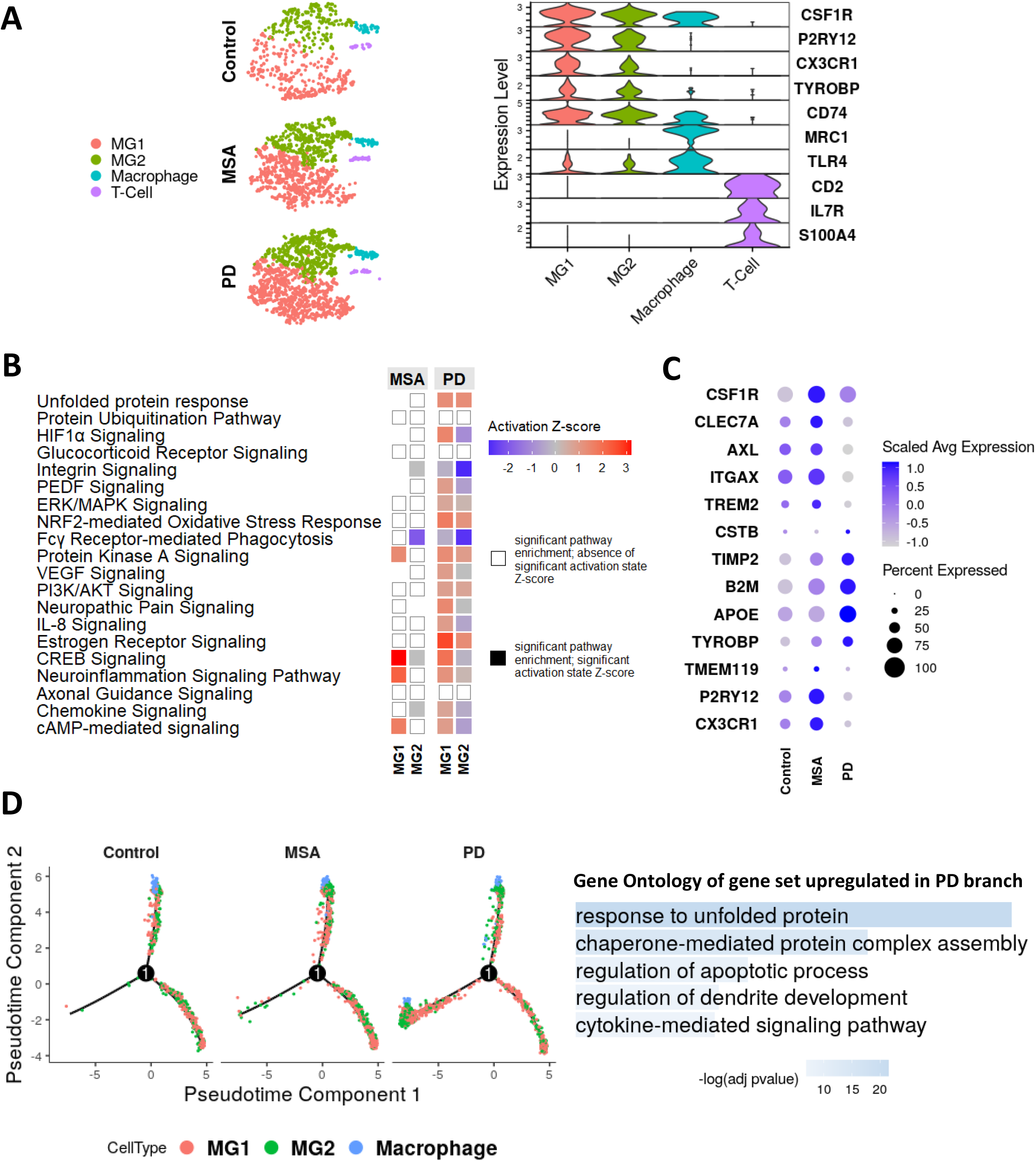
Elevated Response to Unfolded Protein in Activated PD Microglia. **(A)** Microglia population is primarily made up of two clusters -MG1 and MG2 and a Macrophage cluster characterized by expression of MRC1, TLR4 **(B)** Activation zscores of enriched pathways in microglia clusters compared to Control **(C)** Overall expression dot plot of activation genes in microglia, aggregated by disease state, shows APOE dependent activation in PD microglia **(D)** Trajectory reconstruction identifies PD specific branch/state; Genes upregulated in the PD specific branch, compared to other branches, were enriched for unfolded protein response.

### Upregulation of Unfolded Protein Response and Activation of Microglia in PD

Several recent single-cell studies have discussed about the presence of a specific disease associates activated microglia types [14]. We wanted to check the presence of these microglial species in PD and MSA samples. A scaled expression dot-plot of previously identified disease associated microglia (DAM) genes is shown in Figure 4C. We observed upregulation of genes such as TYROBP, APOE, B2M, TIMP2, CSTB and downregulation of P2RY12 and CX3CR1 in PD microglia compared to Control, consistent with previously suggested APOE-dependent / TREM2-independent microglial activation signature [14]. We performed trajectory reconstruction of microglia by considering the top 1100 differentially expressed genes across microglia clusters and disease condition (see Methods section). The branches were labelled as branch-left, branch-up and branch-right, signified by the direction they point to as shown in Figure 4D. The PD specific branching observed here suggests that although PD and MSA are both Synucleinopathies, PD microglia exhibit stark differences from MSA compared to Controls. Genes upregulated in the PD specific branch were functionally enriched (using Enrichr [15, 16]) for Unfolded Protein Response and chemokine-mediated signaling as shown in Figure 4D. Several member genes of UPR -DNAJA1, DNAJB1, HSPA8, HSP90AA1, HSP90AB1, HSPH1, HSPB1 were upregulated in the PD specific branch.

### Differential modulation of pathways in PD and MSA Astrocytes

Astrocytes were the third most abundant cell type, following oligodendrocytes and neurons. We re-clustered astrocytes and identified 3 sub-populations AS1, AS2 and AS3. All clusters expressed AQP4 (Aquaporin 4) and SLC1A3 (aka Excitatory amino acid transporter 1, EAAT1), which are known astrocyte markers. Figure 5A shows t-SNE plot of the three astrocyte clusters in two dimensions and violin plot of genes which are differentially expressed among the astrocyte clusters of Control group. We observed decreased expression of major astrocytic glutamate transporters -SLC1A3 (EAAT1) and SLC1A2 (GLT-1 glutamate transporter 2) in AS2 and AS3 clusters as compared to AS1 cluster. Moreover, GFAP (Glial Fibrillary Acidic Protein) and CD44 were upregulated in AS2 and AS3 astrocyte clusters. This expression pattern indicates that AS1 cluster is likely a quiescent astrocyte population, while AS3 cluster is suspected to be a reactive astrocyte population with the highest upregulation of reactive astrocytic markers GFAP and CD44. Reduced expression of MERTK and MEGF10 in AS3 cluster points to reduced phagocytic capacity of reactive astrocytes [17]. We observed an increased proportion of reactive AS3 population in PD as compared to Control and MSA (Figure 5D), however, no statistical significance was established potentially due to the small sample size (n=3). We then performed differential gene expression analysis of PD versus Control and MSA versus Control nuclei derived from clustering. Subsequent pathway enrichment with IPA revealed downregulation of NRF2-mediated oxidative stress response genes involving reduced expression of chaperone DNAJ heat shock proteins (DNAJ A1/A4/B1/B2/B6/C1/C3) and EIF2AK3 (eukaryotic translation initiation alpha kinase 3) in AS1 and AS2 MSA astrocyte clusters compared to Control group. Genes involved in EIF2 signaling and Oxidative Phosphorylation were upregulated in PD clusters (Figure 5B). Specifically, constituents of the Cytochrome C Oxidase complex such as COX4I1, COX5B, COX6B1, COX6C genes and of the mitochondrial complex I, such as NDUF A3/A4/A6/B4/B10 were upregulated in PD associated astrocytes. A complete list of differentially expressed genes for astrocytes is available in the form of Supplementary Table 4.

**Figure 5.**
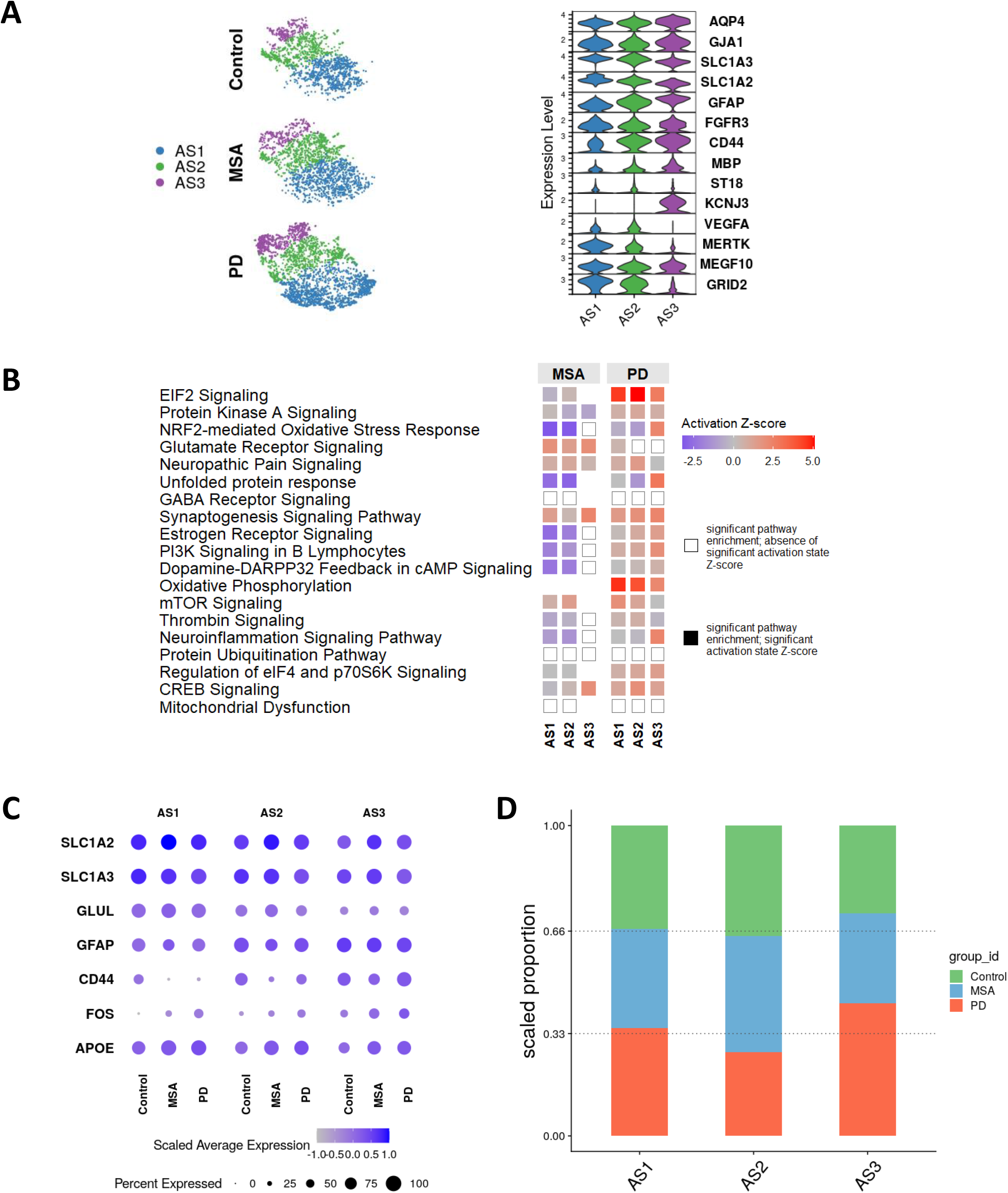
Astrocytic Reactivity. **(A)** Astrocyte clusters AS1, AS2 and AS3 lie on a continuum in 2D t-SNE manifold, ordered according to increasing reactivity accompanied with increasing expression of GFAP and CD44 **(B)** Differential modulation of enriched biological pathways in MSA and PD clusters compared to Control, underscores contrasting altered heterogeneity in PD and MSA **(C)** Scaled average expression of astrocytic reactivity and homeostasis genes, aggregated by disease state **(D)** Reveals trend of increased reactive AS3 astrocytes proportion in PD compared to Control

### Differentially expressed genes in PD cell types are enriched for PD GWAS linked variants

To investigate if altered gene expressions in PD cell-types compared to Control group are linked with PD GWAS variants, we checked the overlap between cell type differentially expressed gene sets and list of genes nearest to significant variants discovered in PD GWAS [18]. The DEG list for each cell type was obtained by merging differentially expressed genes for PD versus Control group cell subtypes. As seen in Figure 6A, the intersection set size for astrocytes, neurons and microglia was 23 (p-value 2.06e-05), 17 (p-value 3.172e-03) and 9 (1.596e-02) genes, respectively. In Figure 6B, we display expression levels of genes which have been consistently implicated in several neurodegenerative diseases, some of which were enriched in PD GWAS variants. The presence of SNCA pathology is known to be concomitant with other important neurodegeneration associated proteins, such as APP and MAPT. We observed higher expression of SNCA in PD microglia and oligodendrocytes and neurons, compared to Control, as shown in Figure 6B. Upregulation in MAPT expression was observed in PD oligodendrocytes, whereas expression of APP was increased in MSA oligodendrocytes compared to Controls.

**Figure 6.**
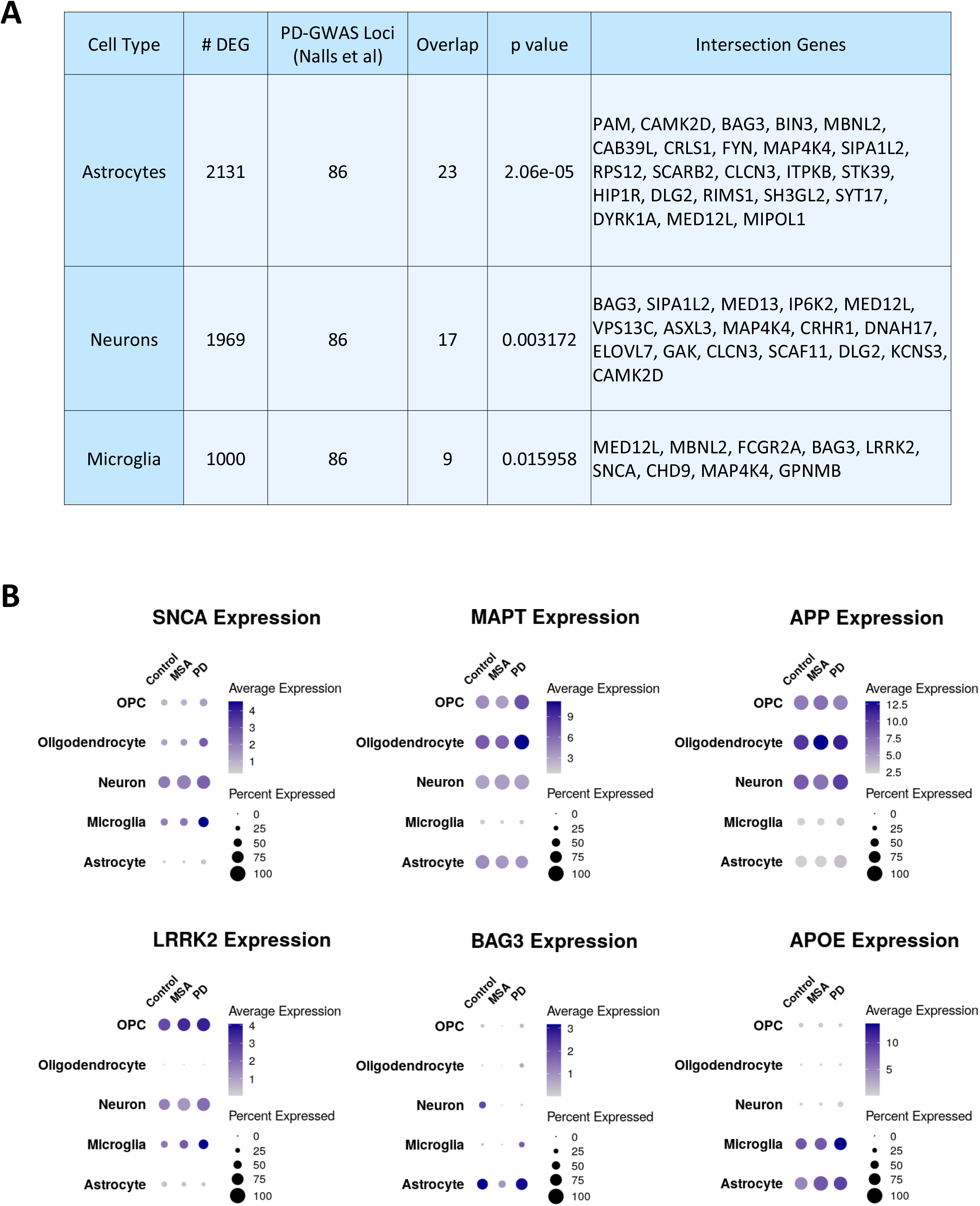
GWAS Enrichment for PD Cell Types. **(A)** Overlap enrichment for differentially expressed genes in PD cell types with significant loci reported in Nalls et al and corresponding intersection genes; Overlap significance is calculated using hypergeometric test **(B)** Expression levels of alpha-synuclein linked genes in different cell types

## Discussion

Understanding of disease biology at the cell-type specific level is crucial in drug discovery and development. In case of major complex neurodegenerative diseases, this cell type specific knowledge has been largely unavailable because of the lack of high throughput single cell profiling techniques for the investigation of postmortem brain samples. Additionally, high sequencing cost has resulted in this area being largely unexplored thus far, with just single PD single-nucleus sequencing study with no publicly available data [7], and none available for MSA. We built this single nucleus expression atlas to molecularly characterize synucleinopathies at cell-type resolution, to broaden biological understanding around α-synuclein regulation and to make the data publicly available through an interactive interface.

Putamen is a part of the forebrain, located at the dorsal striatum and is the outermost structure of the basal ganglia, which is central in the regulation of motor control, cognition, emotions, and learning [19]. This being the first known single cell study from putamen to the best of our knowledge, the cell-type distribution in the human putamen is not well understood. We profiled a total of 87,086 cells from the putamen of 9 subjects and found that oligodendrocytes are the major population (62.88%) in the putamen followed by neurons (17.42%). The abundance of the oligodendrocytes in the putamen suggest that they may be playing a major role in synucleinopathy disease biology. While Agarwal et. al. showed that oligodendrocytes have a significant enrichment for PD GWAS loci [20], MSA has been shown to have α-synuclein containing aggregates in the oligodendroglia fraction [21]. We have comprehensively investigated oligodendrocyte lineage population in our sister study [22]. Consistent with the hypothesis, we found distinct PD and MSA specific changes in the oligodendrocyte fraction [22].

Understanding cellular heterogeneity is not only important to understand the molecular functioning and physiology but also the change in heterogeneity in a disease condition could provide clue for underlying disease mechanisms. We found an overall change in the heterogeneity both in case of PD and MSA (Figure 1C). We found an increased microglial proportion and a decreased OPC proportion in PD compared to Control and MSA group (Figure 1C). An increase in VLMC proportion was also observed both in PD and MSA. Medium spiny neurons are the major inhibitory neuronal cell types in the midbrain. Apart from the established D1 and D2 midbrain medium spiny neuron phenotypes, we detected presence of unresolved ADARB2^+^ SPNs (MSN cluster) expressing both DRD1 and DRD2 receptor genes [23]. These types of SPNs accounted for 6.36% of neurons in the Control group. The distinct alteration of cellular heterogeneity could suggest the existence of different underlying pathological events in PD and MSA. Indeed, there have been reported differences in α-synuclein aggregation in these diseases, both in terms of species of α -synuclein involved and in the regional pattern of aggregates seen in the brain [24]. Validation of these findings would require analysis of a larger sample in the future.

PD and MSA are neurodegenerative disorders presenting with multitude of common symptoms (like parkinsonism, autonomic dysregulation, and progressive ataxia) and shared therapeutic treatment compounds like Levodopa. Given the motor dysfunction in these disorders, neurons play a critical role in their pathologies. Despite of the similarities, we observed contrasting molecular changes in PD and MSA compared to control subjects. D1 and D2 spiny projection neurons make up majority of striatal neurons, however they exhibit differential modulation of enriched biological pathways in PD and MSA (Figure 2B). Dopamine-DARPP32 feedback signaling was activated in PD D1/D2 receptor neurons, whereas it was inhibited in the case of MSA D1/D2 neurons. Moreover, the Brown module of disease correlated genes, identified from WGCNA analysis, had opposite expression pattern in MSA and PD compared to Control neurons. PPI network genes of this module were enriched for dopaminergic synapse, MAPK signaling and mitophagy. These findings, together, suggest contrasting neuronal biology in MSA and PD regardless of the similarities in clinical manifestations, and warrant further studies for comprehensive understanding.

Microglia are the major immune cell type in CNS and have multiple functions from immune surveillance to maintaining synapses and pruning synapses. Recent single cell studies have put microglia into spotlight for neurodegenerative disease by showing the presence of a disease-associated population that could be playing a major role in neuroinflammation leading to a rapid synapse dysfunction [14]. UPR has been shown to play vital role in maintaining synaptic architecture and neuronal signaling [25]. Contrary to PD, absence of upregulated unfolded protein response in MSA microglia, accompanied with neuroinflammation may help explain rapid loss of synapse in MSA. This may point to failure of microglia in responding to α-synuclein aggregates, leading to glial cytoplasmic inclusions (GCIs), which are pathognomonic of MSA [26]. Chronic inflammation is a well-established feature of neurodegenerative diseases [27], which can contribute to dopaminergic neuronal loss [28]. Kim et al. observed that neuron-released α-synuclein promotes activation of LRRK2 in Lipopolysaccharide (LPS) induced rodent microglia, triggering proinflammatory cascade through release of cytokines [29, 30]. Increased LRRK2 mRNA levels accompanied with increased striatal α-synuclein deposits in postmortem PD patients in a previous study [31] suggests that LRRK2 and SNCA might be co-regulated. We observed increased expression of SNCA in neurons, oligodendrocytes, and microglia of PD patients but upregulation of LRRK2 was only observed in neurons and microglia. Therefore, cell specificity seems to be important for the interplay between LRRK2 and SNCA. Both SNCA and LRRK2 have been repeatedly implicated in several PD GWAS studies and APOE is one of the top therapeutic targets for PD [32]. Substantial increase in expression of these genes, combined with PD specific trajectory state, highlights microglia role in PD pathogenesis. These findings should be further explored experimentally to fully characterize these distinct microglia population in PD and MSA and how they contribute to selective pathology.

Increasing number of studies now show that microglia and astrocytes work in tandem [33][16] and the activation of microglia drives the reactivity in astrocytes as well as the other way around. In addition to direct cell-cell interaction, microglia and astrocytes have been shown to interact via secretory molecules. We found that differentially expressed genes in PD and MSA astrocytes match to different regulation of the pathways in PD and MSA astrocytes (Figure 5B), with unfolded protein response and oxidative phosphorylation related gene being upregulated in PD. Transcriptomic profiling of astrocytes enabled identification of three clusters characterized by expression levels of astrocyte reactivity (CD44 / GFAP) and homeostasis (SLC1A2 / SLC1A3) genes. Astrocyte clusters, ordered in increasing level of reactivity – AS1, AS2 and AS3, exhibited differential regulation of genes within different biological pathways. For instance -EIF2 signaling, Protein Kinase A signaling, and oxidative phosphorylation related genes were upregulated in PD whereas genes within the NRF2-mediated oxidative stress response were downregulated in MSA compared to Control. Along with increased proportion of PD microglia, we noted increased proportion of reactive AS3 population in PD which reinforces the idea of interlaced functioning of astrocytes and microglia and possibly activated microglia induced reactive astrocytes.

Overall, this study provides a valuable and unique resource, a transcriptomic atlas at single cell resolution for the putamen region of the human midbrain for two major synucleinopathies: PD and MSA. Even though both PD and MSA are manifestation of the hallmark α-synuclein accumulation, it is interested to see distinct transcriptomic signatures among the two that corroborates the distinct pathology and distinct clinical phenotypes. While our findings require confirmation through analysis of additional patient samples, the PD-and MSA-associated transcriptomic patterns observed among different cell types suggest novel hypotheses for synucleinopathy pathogenesis which remain to be further explored. Implementation of novel proteomics and metabolomic profiling at single cell resolution will provide a more comprehensive understanding of the molecular changes that underpin these two pathologies. We invite researchers to make use of this data and to facilitate this interaction we are providing a web interface to interrogate this dataset.

## Methods

Post-mortem fresh-frozen unfixed human putamen samples were each obtained through partnerships with licensed organizations with completed pre-mortem consent for donation and ethical committee approval for sample acquisition and use. Samples used for single-nucleus RNA sequencing (snRNA-seq) were putamen tissue sections from nine human donors (n =3 per group, mean age in years ± SD: Control, 78.7 ± 9.5; PD, 79.7 ± 5.5; MSA, 65.0 ± 10.6) [22].

### Sample Collection/Nuclei Sorting/Suspension/Barcoding/Library Preparation

Sample procurement, nuclei sorting, and library preparation is described in Teeple et. al. [22].

### QC, Integration, Clustering and Cell Type Annotation

As part of quality control, we filtered nuclei with aberrant gene capture (nFeature_RNA) and mitochondrial gene percentage (percent.mt) to reduce multiplets and low-quality nuclei in downstream analysis. Thresholds for these measures were inferred from violin plots as shown in Figure 1A and are as follows: 200 < nFeature_RNA < 9000 and percent.mt < 5. Following quality control, gene counts were normalized using the NormalizeData() function (Seurat v3.2 [34]; R version 3.6.1) with scale factor as 10,000 and LogNormalize() method. With the top 2000 most variable features from the FindVariableFeatures() function, samples were then integrated using Seurat’s standard integration workflow (FindIntegrationAnchors() and IntegrateData() functions). We clustered nuclei with the Louvain algorithm (FindClusters() Seurat function) using the first 7 principal components. After experimentation, the resolution hyperparameter was set to 0.2, based on visual separation among the clusters. One cluster with exceptionally low gene capture and three sparsely populated clusters expressing multiple cell type markers were filtered out and not included in downstream analyses.

### Cell Subtype Clustering

Neuronal subset was reintegrated using 1000 anchor features and clustered with first 12 principal components and 0.1 resolution parameter. Out of the 10 obtained clusters, one cluster expressed markers for both oligodendrocyte as well as neuron, therefore this cluster was dropped from further analyses. Three clusters which had particularly low nuclei count, were also excluded from downstream analyses. The remainder clusters were annotated by exploring differentially expressed genes in the Control group using FindMarkers() function in Seurat.

We considered 1000 anchor features to reintegrate astrocyte fraction and a resolution parameter of 0.2 along with the first 7 principal components to re cluster astrocytes. Of the five derived clusters, one cluster was populated primarily by nuclei from single PD sample (PD4), while another cluster expressed multiple cell type markers for oligodendrocyte as well as astrocyte. We excluded both these clusters from further analyses.

Following neurons and astrocytes, the microglia subset was subsequently reintegrated using 1000 anchor features and re clustered with first 4 principal components and 0.1 resolution parameter. A sparsely populated cluster (106 nuclei) expressed T cell markers (IL7R, CD2, S100A4) with almost no expression of CSF1R. Suspecting the origin of this cluster to be blood contamination of samples, we excluded this from downstream analysis.

### Pathway Enrichment Analysis

For each of the sub-population of a given cell-type, differential gene expression for PD versus Control and MSA versus Control was performed using the FindMarkers() function in Seurat (v3.2) and the MAST R package [35]. Canonical pathway regulation analysis was generated from the result of differential gene expression analysis, through the use of IPA software (QIAGEN Inc., https://www.qiagenbioinformatics.com/products/ingenuity-pathway-analysis). We used adjusted p-value cutoff of 0.05 and log fold change cutoff of 0.25 (fold change cutoff of ∼ 1.28, ∼ e^0.25^) for core expression analysis in IPA. The Z-score reported by IPA assesses the match of observed and predicted up/down regulation patterns within gene sets [36]. Missing Z-scores represent absence of a statistically significant activation state of the regulator.

### Pseudo-time Trajectory Reconstruction Analysis

For performing trajectory analysis using DDRTree algorithm [37] in Monocle 2 [8], we considered genes that were differentially expressed across cell type and condition while removing the effects of individual samples, determined by the function differentialGeneTest(). This allows for the reconstructed trajectory to capture heterogeneity as well as activation information. The number of genes (sorted by p-value) used for ordering cells on trajectory was treated as a hyperparameter and optimized to reduce sparsely populated branches.

### Weighted Gene Co-expression Network Analysis (WGCNA)

WGCNA affords identification of gene sets with similar expression patterns across different nuclei. We used the WGCNA R package in R [38]. We used the cell x gene expression data from Seurat object, after correcting for batch effects using SCTransform function from Seurat with vars.to.regress parameter set to day of sample sequencing, and performed hierarchical clustering, dynamic tree cutting and subsequently identified 12 co-regulated gene modules. Furthermore, according to WGCNA protocol, we calculated Pearson correlations between module eigen vectors and one-hot encoded disease condition to check if any gene modules are related to disease condition. Positive correlation indicates higher expression of module genes, while negative correlation indicates reduced expression; and the absolute value of the correlation coefficient signifies the strength of association between module gene set and disease state. After identifying modules that are highly associated with disease condition, we used the module genes to generate tissue specific (putamen) functional protein-protein interaction network (PPI) using NetworkAnalyst [39] and DifferentialNet database [40]. For further investigation, network node genes were then analyzed for their association with biological pathways. Proteins from minimum PPI network were probed for overlap with genes within 100kb regions of PD GWAS variants (association p value < 1e-5) from the 2019 study by Nalls et. al. [18]. Statistical significance for overlap enrichment was examined using phyper() hypergeometric test function in R. Intersection overlap Venn diagram was generated using InteractiVenn [41].

### Cell Type PD GWAS Enrichment Analysis

For each PD cell type we merged differentially expressed genes identified as sub cell type resolution (compared to corresponding Control group), followed by overlap enrichment with genes closets to significant PD GWAS variants. As mentioned above, statistical significance for overlap was established using hypergeometric test. This analysis enabled is to inspect if gene expression variations in a particular cell type are exceedingly enriched for PD GWAS [18]. To check if gene expression changes in a cell type are associated with PD GWAS variants.

## DATA AND CODE AVAILABILITY

Single nuclei atlas (powered by cellxgene: https://chanzuckerberg.github.io/cellxgene/) is available at <link>. For a list of all supported features of the interface, visit https://chanzuckerberg.github.io/cellxgene/posts/gallery.

R markdown code files of the analysis will be made available at the time of publication.

Seurat v3 toolkit for single cell genomics:

https://satijalab.org/seurat/

WGCNA package: Installation and documentation

https://horvath.genetics.ucla.edu/html/CoexpressionNetwork/Rpackages/WGCNA/

NetworkAnalyst online interface:

https://www.networkanalyst.ca/

Enrichr: Gene list enrichment analysis tool:

https://amp.pharm.mssm.edu/Enrichr/

InteractiVenn: Analysis of sets through Venn diagrams

http://www.interactivenn.net/

## ACKNOWLEDGMENTS

We thank Dr. Srinivas Shankara for his critical review and insightful feedback on this paper.

## AUTHOR CONTRIBUTIONS

All authors have each made substantial contributions to study concept, design, and implementation; drafting and critically revising the manuscript for important intellectual content; and provided final approval of the submitted manuscript version.

---

**Supplementary Figure 1.**
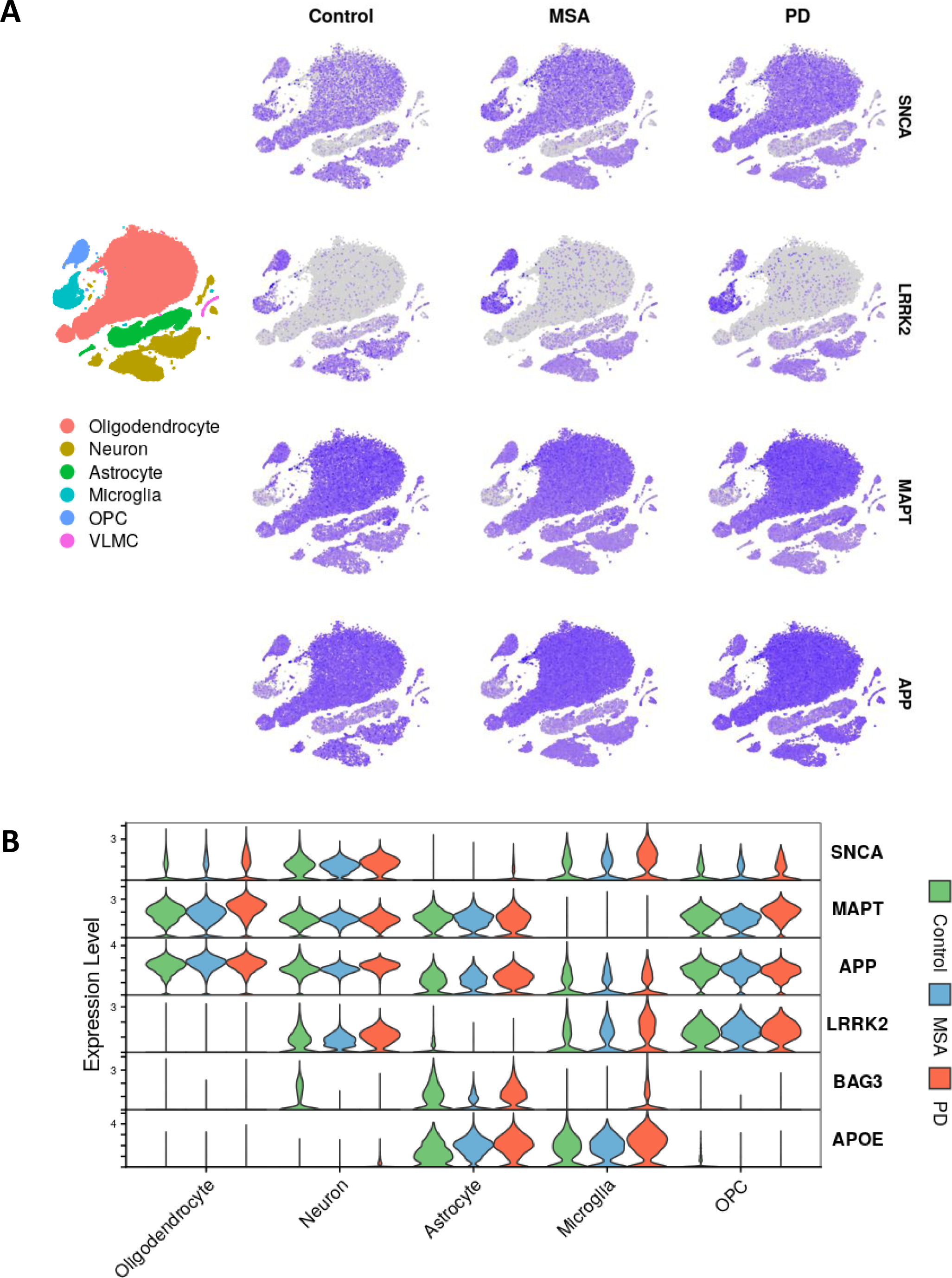
Gene Expression Profiles. **(A)** 2D t-SNE plot of nuclei colored by gene expression levels of SNCA, MAPT and APP **(B)** Violin expression plot of SNCA, MAPT, APP, LRRK2, BAG3 and APOE in different cell types

**Supplementary Table 1.** Canonical marker genes for major brain cell types.

| Cell Type | Marker Genes |
| --- | --- |
| Oligodendrocytes | BCAS1, OPALIN, HAPLN2 |
| OPCs | PDGFRA, VCAN |
| Neurons | RBFOX3 |
| Astrocytes | AQP4, GFAP |
| Microglia | CSF1R, CD74 |
| VLMC | DCN, COLEC12, PDGFRB |

**Supplementary Table 2** | **Differential gene expression data of PD and MSA neuronal sub-types against control**. Data for each of the PD and MSA neuronal subtypes, for is organized in separate tabs of the worksheet.

**Supplementary Table 3** | **Differential gene expression data of PD and MSA microglial sub-clusters against control**. Data for each of the PD and MSA microglial clusters is organized in separate tabs of the worksheet.

**Supplementary Table 4** | **Differential gene expression data of PD and MSA astrocyte sub-clusters against control**. Data for each of the PD and MSA astrocyte clusters is organized in separate tabs of the worksheet.

**Supplementary Table 5** | **List of co-expressed gene modules in Neurons across disease states**

**Supplementary Table 6.** Broad cell type nuclei counts aggregated by putamen sample.

| Sample | Oligodendrocyte | Neuron | Astrocyte | Microglia | OPC | VLMC |
| --- | --- | --- | --- | --- | --- | --- |
| CTR1 | 3599 | 411 | 308 | 235 | 223 | 4 |
| CTR3 | 3078 | 1634 | 1082 | 304 | 478 | 58 |
| CTR8 | 7238 | 1436 | 554 | 315 | 417 | 42 |
| MSA10 | 6125 | 2757 | 761 | 617 | 265 | 144 |
| MSA5 | 7784 | 501 | 580 | 525 | 412 | 15 |
| MSA7 | 5472 | 3136 | 949 | 366 | 557 | 99 |
| PD2 | 9049 | 126 | 542 | 532 | 259 | 9 |
| PD4 | 3940 | 3598 | 2421 | 979 | 271 | 155 |
| PD9 | 6324 | 976 | 1160 | 491 | 270 | 80 |

**Supplementary Table 7.** Neuronal sub type nuclei counts aggregated by putamen sample.

| Sample | D1MSN | D2MSN | MSN | GABA1 | GABA2 |
| --- | --- | --- | --- | --- | --- |
| CTR1 | 109 | 210 | 18 | 17 | 7 |
| CTR3 | 597 | 631 | 84 | 83 | 10 |
| CTR8 | 541 | 518 | 87 | 32 | 25 |
| MSA10 | 923 | 1283 | 103 | 79 | 57 |
| MSA5 | 186 | 186 | 19 | 25 | 11 |
| MSA7 | 1066 | 1340 | 189 | 71 | 73 |
| PD2 | 26 | 51 | 11 | 7 | 2 |
| PD4 | 1402 | 1237 | 232 | 133 | 200 |
| PD9 | 316 | 415 | 61 | 26 | 23 |

**Supplementary Table 8.** Microglial sub type nuclei counts aggregated by putamen sample.

| Sample | MG1 | MG2 | Macrophage | T-Cell |
| --- | --- | --- | --- | --- |
| CTR1 | 120 | 68 | 6 | 13 |
| CTR3 | 114 | 100 | 19 | 3 |
| CTR8 | 149 | 90 | 15 | 5 |
| MSA10 | 249 | 215 | 33 | 34 |
| MSA5 | 281 | 148 | 10 | 5 |
| MSA7 | 202 | 97 | 17 | 8 |
| PD2 | 290 | 154 | 12 | 4 |
| PD4 | 534 | 327 | 29 | 11 |
| PD9 | 216 | 147 | 32 | 23 |

**Supplementary Table 9.** Astrocyte sub clusters nuclei counts aggregated by putamen sample.

| **Sample** | **AS1** | **AS2** | **AS3** |
| --- | --- | --- | --- |
| CTR1 | 108 | 113 | 65 |
| CTR3 | 541 | 388 | 48 |
| CTR8 | 230 | 138 | 113 |
| MSA10 | 365 | 208 | 106 |
| MSA5 | 187 | 168 | 113 |
| MSA7 | 430 | 400 | 49 |
| PD2 | 93 | 291 | 91 |
| PD4 | 1252 | 252 | 277 |
| PD9 | 354 | 357 | 263 |

